# The Human Male Mammary Gland has Similar Epithelial Populations to Female but Distinct Composition and Transcriptional Properties

**DOI:** 10.64898/2026.03.27.714915

**Authors:** M-Iliana Ibañez-Rios, Syed Mohammed Musheer Aalam, Megan L. Ritting, Ashlyn Jore, Keyur Chaludia, Chitra Priya Emperumal, James W. Jakub, Sarah A. McLaughlin, Amy C. Degnim, Judy C. Boughey, Fergus J. Couch, Siddhartha Yadav, Anguraj Sadanandam, Mark E. Sherman, Derek Radisky, David J.H.F. Knapp, Nagarajan Kannan

## Abstract

The normal adult male breast has not been characterized at single-cell resolution, leaving the cellular basis of male breast cancer (MBC) biology undefined. Here we present an integrated single-cell RNA sequencing atlas of the adult human breast comprising 174,471 cells from 17 donors (3 male, 14 female), including 18,117 male-derived cells. This revealed that the male breast retains all three epithelial populations, basal (BC), luminal progenitor (LP), and luminal committed cells (LC), but with an increase in LC at the expense of BC and LP across all three male donors. Male LC were distinguished from female by elevated *ESR1* and *PGR* mRNA, enrichment of RNA processing and ribosome biogenesis programs, reduced inflammatory cytokine and growth factor signaling, elevated estradiol gene set enrichment scores, and higher inferred activity of developmental patterning transcription factors. This pattern was observed across differential expression, gene ontology, ligand profiling, and regulon-based analyses, and was not restricted to sex chromosome-linked gene expression. This is consistent with the near-universal estrogen receptor (ER) positivity that characterizes MBC clinically. This atlas provides the first cellular and transcriptional reference for the normal male breast and a resource for investigating sex differences in mammary biology, germline susceptibility variant interpretation, and modeling breast malignancies.

## INTRODUCTION

The mammary gland is a sexually dimorphic tissue system which is one of the defining characteristics of mammals. In human, early breast development proceeds similarly in both sexes; however, divergent hormonal environments at puberty produce distinct outcomes. Estrogen and progesterone drive ductal elongation and lobuloalveolar differentiation in females, while high circulating androgens suppress epithelial expansion in males, yielding a rudimentary ductal system largely devoid of terminal duct lobular units.^1–3^ In most mammals including common laboratory models such as mice, androgen exposure results in active regression of the mammary bud in development, while in humans a rudimentary ductal system is retained, though one largely devoid of terminal duct lobular units.^1,2^ In humans, this residual ductal system retains some capacity for hormone-responsive remodeling as is illustrated by gynecomastia, in which altered androgen-to-estrogen ratios produce benign ductal proliferation, and by documented lactation induction in biological males receiving lactogenic hormonal therapy.^4–6^ These observations indicate that hormone-responsive epithelial cells persist in adult human male breast tissue. While the identity, abundance, and transcriptional characteristics of the major epithelial populations of the human female breast have been well described through prospective isolation and single-cell profiling,^7,8^ no equivalent characterization of the male breast has been reported.

Clinically, male breast cancer (MBC) is distinguished from female breast cancer by its near-universal ER positivity, reported in up to >90% of cases across population-based series. This prevalence substantially exceeds the approximately 75–80% observed in female breast cancer, rendering MBC one of the most hormonally homogeneous of all breast cancer subtypes. MBC also presents more frequently at higher histological grade and carries a distinct germline susceptibility landscape relative to female disease.^9–15^ Given its rarity, MBC has limited prospective clinical trial data to guide treatment;^16^ management decisions rely on extrapolation from small retrospective series and from trials conducted exclusively in women.^17^ While single-cell transcriptomic profiling has been applied to male breast tumors,^8^ the normal, non-cancerous male mammary epithelium has not previously been characterized at single-cell resolution, leaving the cellular baseline against which tumor-associated transcriptional changes must be interpreted undefined. Despite this clinical distinctiveness, the cellular context in which these epidemiological and genetic differences arise, that is, the composition and molecular state of the normal male breast epithelium, has not been defined. This represents a gap in the foundational biology needed to interpret MBC risk, model the disease, or contextualize susceptibility variants.

The human mammary epithelium is organized as a hierarchical bilayer in which a basal/myoepithelial compartment encloses an inner luminal layer comprising two distinguishable populations: luminal progenitor (LP) cells, defined as Lin⁻CD49f⁺EpCAM⁺, and luminal committed (LC) cells, defined as Lin⁻CD49f^low/−^EpCAM⁺.^18–21^ LC and LP are also referred to as the luminal hormone-sensing (LHS) cell, the luminal adaptive secretory precursor (LASP) cell in single cell omics studies.^22^ These populations differ in their hormone sensitivities, proliferative outputs, and cancer subtype associations. In the female breast, LP carry transcriptional programs associated with basal-like and *BRCA1*-linked breast cancers, while LC maintain programs resembling ER⁺ luminal disease.^19,23^ These associations are derived from transcriptional comparisons between sorted normal populations and tumor profiles and, given the known plasticity of epithelial cells, are best interpreted as candidate rather than established origin relationships.^19,24–27^ The capacity to prospectively isolate these populations and characterize them at the transcriptomic level has substantially advanced understanding of normal mammary biology and cancer susceptibility in women.^19,28,29^ Despite the existence of a morphologically recognizable, hormone-sensitive mammary epithelium in males,^30^ comparable single-cell characterization has not been reported. Understanding the composition and function of the male mammary gland cells, is critical both for understanding male breast cancer, and as a counterpoint to females, for elucidating the effects of hormonal environment in human mammary tissue more generally, differences that are not well captured by animal models given the developmental differences in mammary gland retention.

To define the cellular composition and transcriptional identity of the adult human male breast, we integrated scRNA-seq data from male and female donors to construct a first sex-stratified single cell atlas of normal mammary cells. This analysis reveals a LC-dominant epithelial architecture in the male breast, characterized by elevated hormone receptor expression, attenuated cytokine signaling, and sex-biased transcription factor activity, providing the first normal tissue reference for investigating hormonal regulation of the male mammary gland and the cellular origins of male breast cancer.

## RESULTS

### Male Breast Epithelium Is Sparse, Variable Across Donors, and Proliferates More Slowly *In Vitro*

We assembled a cohort of 8 adult male donors (mean age 60.5 years; range 32–77), alongside 27 adult female donors (mean age 47.2 years; range 32–68) undergoing elective surgery or autopsy at Mayo Clinic (Figure 1A). Cryopreserved tissue organoids were thawed and dissociated into single cells using methods established for female breast tissue^18^ and applied without modification to male samples. Lineage depletion was applied to exclude endothelial (CD31⁺), stromal (CD34⁺CD31⁺), and hematopoietic (CD45⁺) cells; viable Lin⁻ (CD31⁻CD34⁻CD45⁻) DAPI⁻ events were classified into phenotype-based compartments by EpCAM and CD49f co-staining: in female donors, three fractions were resolved, LC (EpCAM^hi^CD49f^low^), LP (EpCAM⁺CD49f⁺), BC (EpCAM^low^CD49f^hi^); in male donors, the atypical EpCAM/CD49f distributions described below precluded consistent resolution of three discrete fractions, and populations are referred to throughout as luminal-like (EpCAM^hi^CD49f^low/+^) and basal-like (EpCAM^low^CD49f^hi^) to reflect their phenotypic rather than confirmed lineage identity.^18^ Non-epithelial fractions were excluded from the primary epithelial analysis; stromal populations were retained and are characterized alongside epithelial cells in the integrated atlas described in the following section, though a dedicated investigation of sex differences in the male breast stromal compartment remains an important future direction.

**Figure 1.**
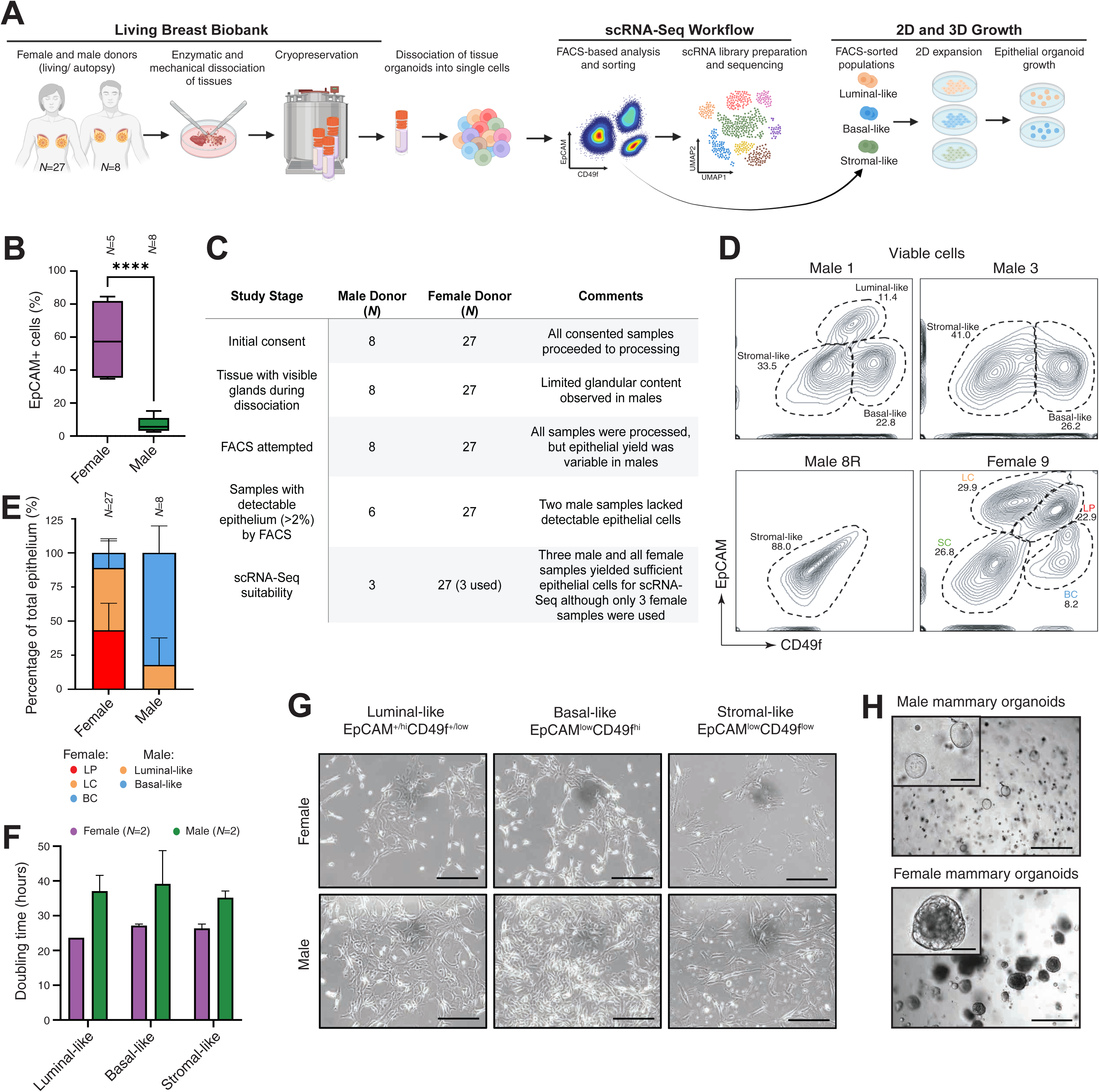
Characterization of male mammary tissue, epithelial content, and *ex vivo* cellular properties. (A) Schematic of study design and workflow, from tissue collection of male (*N*=8) and female (*N*=27) breast tissues through the Mayo Clinic Living Breast Biobank through organoid dissociation, cryopreservation, FACS-based analysis and sorting, scRNA-Seq library generation, and 2D/3D culture assays. (B) Violin plots comparing EpCAM⁺ cell percentage among DAPI⁻ events between female and male donors. (****p<0.0001, Mann–Whitney U test). (C) Summary table showing contrasting outcomes at each processing stage for male and female donors. (D) Representative FACS contour plots of EpCAM vs. CD49f from three male donors (1, 3, 8R) and 1 female donor (9), illustrating inter-donor variability in epithelial phenotype among Lin⁻ (CD31⁻CD34⁻CD45⁻) DAPI⁻ viable cells. Gate percentages indicate fractions of cells within luminal progenitor-like, lumin al-like, basal-like, and stromal-like regions. (E) Female and male stacked bar chart (% ± SD) showing proportional representation of luminal progenitor-like, luminal-like, basal-like, and stromal-like fractions across male and female donors. (F) Bar graph of mean doubling times (hours ± SD) for sorted luminal-like, basal-like, and stromal-like populations from female and male donors under identical culture conditions. (G) Representative images of 2D expanded sorted luminal-like, basal-like, and stromal-like populations from female and male donors under identical culture conditions. Scale bars = 200 µm. (H) Representative images of 3D organoids from female and male donors under identical culture conditions. Scale bars = 200 µm. Inset scale bars = 50 µm.

Epithelial content, measured as the fraction of EpCAM⁺ cells among DAPI⁻ events, was significantly lower in male samples compared to female samples (Figure 1B; 20-70% in female vs 0-15% in male; p<0.0001). Two of eight male donors fell below a threshold of detectable epithelium (>2% EpCAM⁺), and only three yielded sufficient cells for scRNA-Seq library generation, whereas all female samples maintained sufficient epithelial yield for scRNA-Seq, of whom three were selected for library generation (Figure 1C). Representative FACS profiles from male donors illustrate the pronounced inter-donor unpredictability of male mammary epithelial phenotype (Figure 1D). Unlike female donors, in whom EpCAM/CD49f distributions consistently resolved three discrete epithelial fractions, luminal progenitor-like, luminal-like, and basal-like, male donors showed a spectrum of atypical profiles: near-complete absence of detectable epithelium in some donors, collapse of the three expected fractions to two in others, and markedly altered gate boundaries and population proportions across the cohort. These findings indicate that standard FACS phenotyping criteria developed and validated for female mammary tissue cannot be applied with equivalent confidence to male donors, and that surface phenotype alone is an unreliable basis for epithelial subtype classification in the male breast (Figure 1D-E). Accordingly, FACS-defined fractions in male tissue are presented throughout as phenotype-based compartments rather than confirmed lineage assignments; the transcriptomic resolution of epithelial from stromal identity in the scRNA-seq data, described in the following section, provides the more reliable basis for lineage classification in male tissue. This variability likely reflects multiple factors including sex, donor age, body mass index, and hormonal history, though the present dataset is not powered to dissect these contributions.

Sorted populations from male donors across luminal-like, basal-like, and stromal-like fractions retained replicative competence, though they showed longer doubling times than female counterparts under identical culture conditions (Figure 1F-G). Whether this reflects an intrinsic difference in proliferative capacity, a differential response to culture conditions optimized for female mammary cells, or both, cannot be determined from these data alone; the finding is however consistent with the reduced proliferative gene programs observed independently in uncultured male LC by scRNA-seq, described below. Male-derived mammary cells seeded at clonal density in 3D Matrigel generated organoid structures, though outgrowth was less robust than that observed under identical conditions with female-derived cells. Consistent with the reduced growth kinetics, female-derived cells typically formed well-defined organoid structures in less than 7 days of culture, whereas male-derived cultures required closer to 3 weeks to generate comparable structures. Once established, however, the male organoids could be maintained in culture for more than 6 weeks. These data nonetheless establish proof-of-concept that normal male mammary cells retain the capacity for self-organized three-dimensional growth, providing a tractable experimental platform for future functional investigation of male mammary epithelial biology (Figure 1H). These observations establish the practical and biological context for the single-cell analysis: male breast tissue yields fewer epithelial cells with greater variability across donors than female tissue processed by the same methods.

### An Integrated Single-Cell Atlas Reveals Distinct Epithelial Composition in the Male Breast

We generated scRNA-seq profiles from FACS-sorted viable single cells isolated from three normal, non-cancerous male donors and integrated them with in-house data from three female donors and publicly available data from eleven additional female donors,^7^ yielding an atlas of 174,471 cells from 17 donors (Figure 2A). The three male donors contributing scRNA-seq data had a mean age of 66 years (range 56-77); the three in-house female donors had a mean age of 41 years (range 37-46). This age difference between the sequenced male and female cohorts should be considered when interpreting transcriptional differences, particularly those involving hormone receptor expression, which is known to vary with age and menopausal status in female breast tissue. To maximize epithelial representation, the male donor with the highest epithelial yield was sorted as both a total viable single-cell fraction and an EpCAM⁺-enriched viable fraction, generating two complementary scRNA-seq libraries from a single donor; both datasets were retained in the integrated atlas. UMAP visualization revealed coherent clustering by cell type across both sexes, with basal cells (BC), LP cells, LC cells, fibroblasts, smooth muscle cells (SM), endothelial cells, and pericytes resolved as discrete communities (Figure 2B). Cell identities were confirmed by canonical marker gene expression (data not shown). This indicates that the principal transcriptionally defined cell populations of the mammary gland are conserved in the male breast, though transcriptional similarity does not preclude functionally important differences in activity. Notably, the male breast is a ductal-only system lacking terminal duct lobular units; however, LP have been shown to reside in both ductal and lobular compartments of the female breast,^31^ suggesting that the LP population identified here in male tissue is not contingent on lobular architecture for its existence or maintenance.

**Figure 2.**
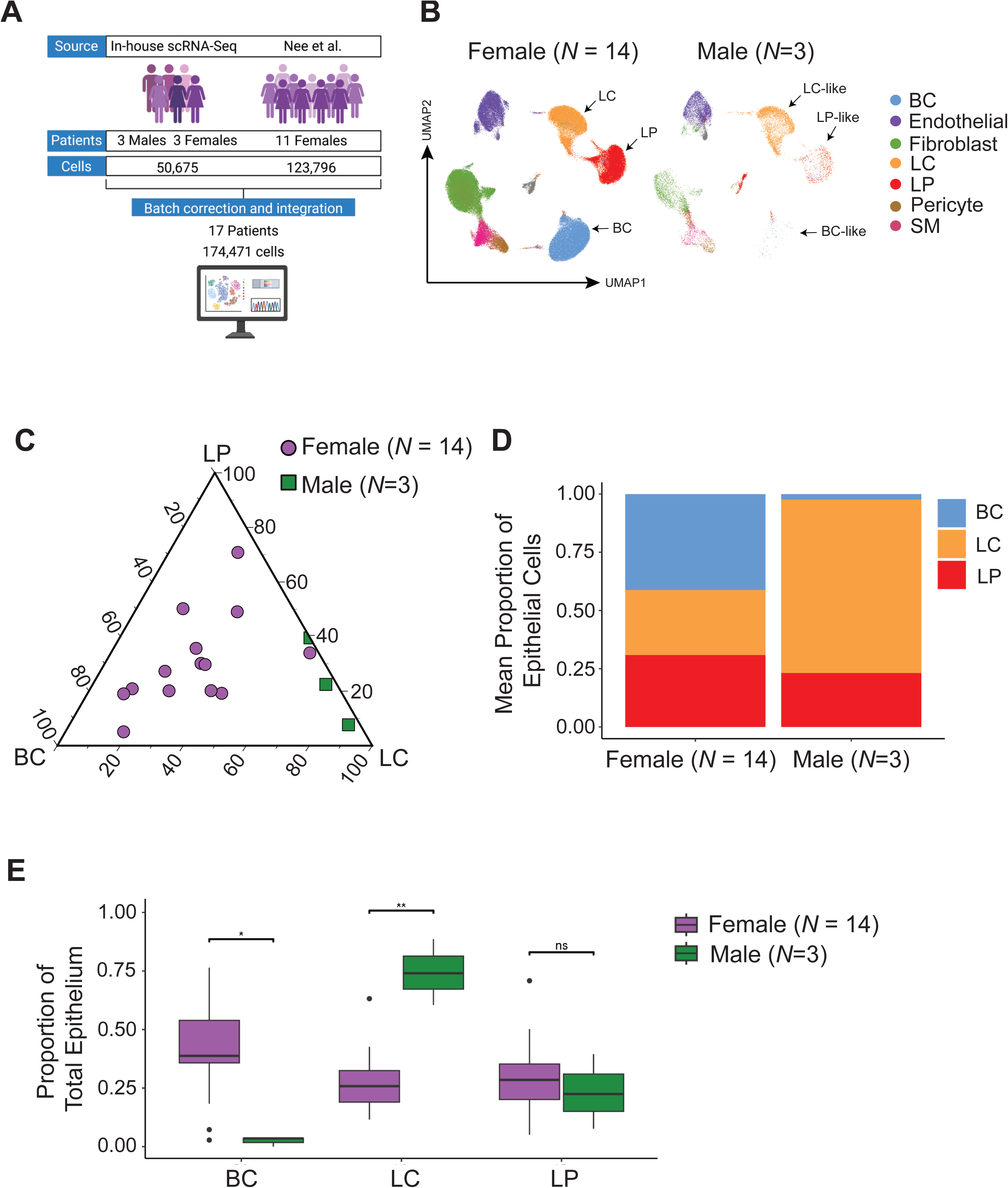
Sex-stratified single-cell atlas of the human breast discovers distinct epithelial subtype proportions in male donors. (A) Atlas construction overview. In-house scRNA-Seq data from 3 male and 3 female donors (50,675 cells) were integrated with publicly available data from 11 female donors (Nee et al.; 123,796 cells) following batch correction, yielding an atlas of 174,471 cells from 17 donors. (B) UMAP visualizations of the integrated atlas stratified by sex (Female: *N*=14; Male: *N*=3). Cell types were inferred based on cluster marker expression profile (C) Ternary plot of BC, LP, and LC fractional composition per donor. Female donors (purple circles) are distributed across the ternary space; male donors (green squares) fall near the LC vertex with markedly reduced BC fractions, in a region not occupied by any female donor. (D) Stacked bar plots of mean epithelial subtype proportions (BC, LC, LP) for female (*N*=14) and male (*N*=3) cohorts, showing higher mean LC proportion and lower mean BC proportion in male donors. (E) Boxplots comparing BC, LC, and LP proportions between female (*N*=14) and male (*N*=3) donors. BC proportion is significantly lower in males (*p<0.05); LC proportion is significantly higher (**p<0.01); LP proportions are not significantly different (ns). Statistical test: Wilcoxon rank-sum.

The most notable feature of male breast epithelial composition was a pronounced shift in the relative abundance of epithelial subtypes. Ternary compositional analysis showed that all three male donors fell near the LC vertex of the BC–LP–LC space, whereas female donors were distributed across a broader region (Figure 2C). Mean LC proportion was higher in males, while mean BC proportion was lower, as shown by compositional bar plots (Figure 2D). Statistical comparison confirmed a significant reduction in BC proportion (p<0.05) and a significant elevation in LC proportion (p<0.01) in males while LP proportions did not differ significantly between groups (Figure 2E).

These scRNA-seq-derived compositional findings appear discordant with the FACS-based subtype profiles shown in Figure 1E, in which male donors show a pronounced basal-like fraction and near-absence of luminal progenitor-like cells by surface phenotype. This discrepancy is resolved at the transcriptional level: within the EpCAM^low^CD49f^hi^ gate used to define the basal-like fraction by FACS, scRNA-seq identified a substantial subpopulation expressing the perivascular marker *RGS5* rather than the canonical basal epithelial markers *KRT5* and *KRT14* (data not shown). In the male breast, where epithelial content is markedly reduced relative to female tissue, this gate is susceptible to contamination by stromal elements, particularly pericytes and smooth muscle cells that share overlapping surface phenotypes with basal epithelial cells but are transcriptionally distinct. The LC-dominant composition documented by scRNA-seq therefore reflects the epithelial architecture of the male breast more accurately than surface phenotype-based classification alone. The flow cytometric data from the broader eight-donor cohort are used to corroborate the LC-enriched luminal-like compartment specifically, the gate least susceptible to stromal contamination, rather than as independent confirmation of basal cell depletion, for which the scRNA-seq data are the primary evidence. Given the small size of the male sequenced cohort (*N*=3), these compositional findings should be understood as reproducible preliminary observations grounded in the broader eight-donor FACS dataset, and independently confirmed in that larger cohort, rather than as definitive quantitative estimates.

### Male LC Are Transcriptionally Distinguished by Elevated RNA Processing Programs and Reduced Cytokine Activity

To ensure the robustness of gene expression changes and eliminate the confounding effects of high and variable cell numbers, we opted for a pseudobulk approach to differential gene and program expression changes. Differential expression analysis between male and female LC identified two broad gene expression patterns stratified by sex (Figure 3A). Complete differential expression results are available upon request and will be included in the final version of the manuscript. Female LC showed higher expression of X-chromosome transcripts (*XIST*, *TSIX*, *KLHL13*), chemokines (*CXCL17*), and transcriptional regulators (*SIX1*, *HOXC10*, *CREB3L1*, *NFKBIE*). Male LC showed higher expression of neurotrophin receptor (*GFRA1*), neuropeptide-related genes (*NPY1R*, *NPY5R*, *NDN*), ion channels (*KCND2*, *KCNK15*, *KCNF1*), fatty acid oxidation genes (*ACADL*, *ACOX2*), protocadherins (*PCDHB2*, *PCDHA10*), Y-chromosome transcripts (*RPS4Y1*, *EIF1AY*, *DDX3Y*), and *CD109* (Figure 3A–B). Gene Ontology analysis of upregulated genes in male LC identified significant enrichment of RNA processing, ribosome biogenesis, and rRNA/tRNA metabolic process terms (Figure 3C), alongside RNA binding and methyltransferase activity at the molecular function level (Figure 3E). These enrichments were not affected by excluding Y-chromosome encoded genes (data not shown). Genes with lower expression in male LC were enriched for immune system process, adaptive immune response, leukocyte activation, and antimicrobial response terms (Figure 3D), and for signaling receptor activity, cytokine activity, and hormone activity at the molecular function level (Figure 3F). These patterns suggest a translationally poised state with limited signaling engagement, suggesting epithelial maintenance rather than active tissue remodeling.

**Figure 3.**
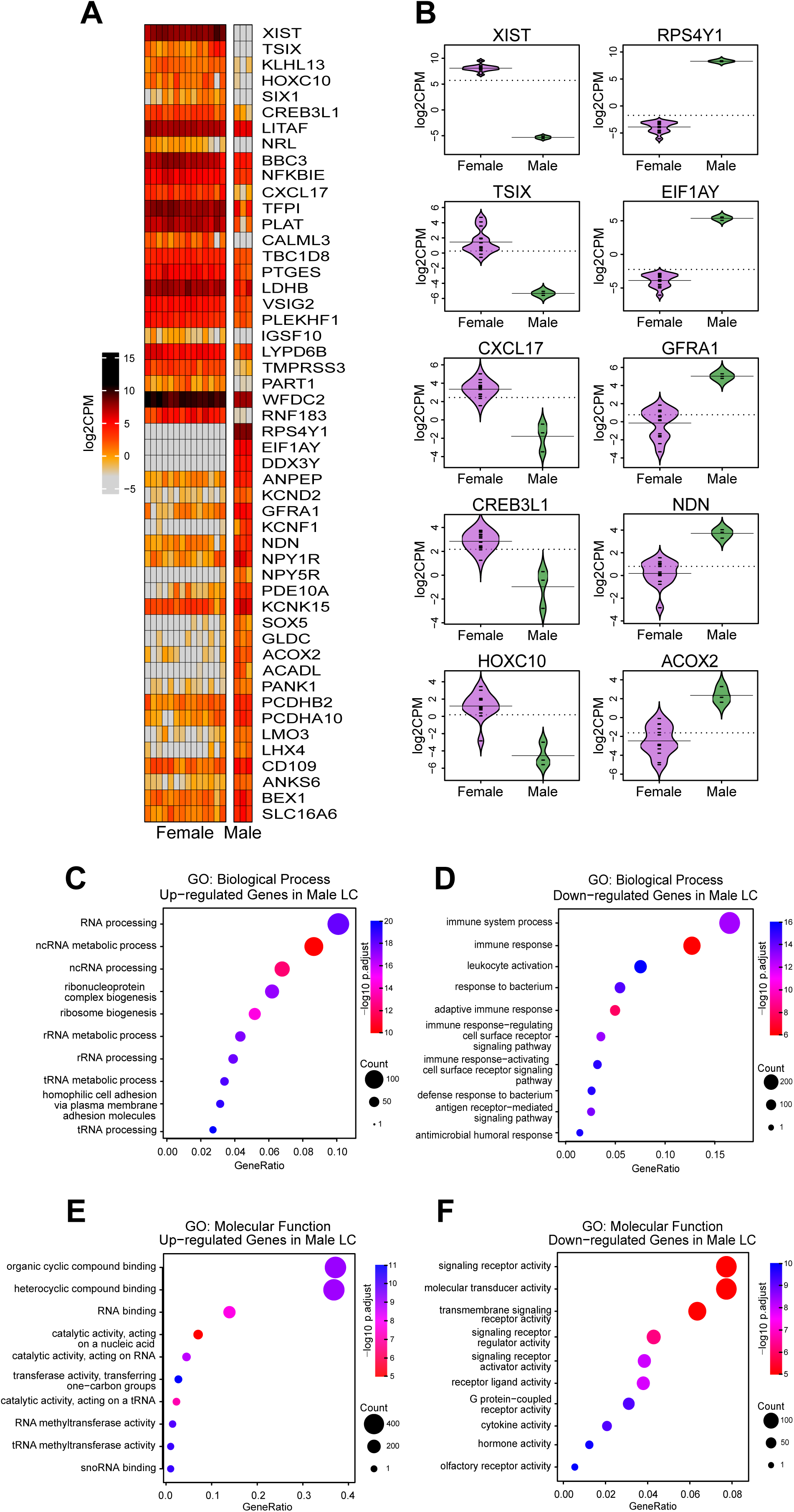
Differential gene expression between male and female LC identifies sex-associated differences in RNA processing and immune-related gene programs. (A) Heatmap of per-donor log2CPM-normalized expression for 50 differentially expressed genes between male and female luminal cells (LC), comprising the top 25 upregulated and top 25 downregulated genes in male versus female LC. Differential expression was assessed by pseudobulk limma-voom (TMM normalization, Empirical Bayes moderation, Benjamini-Hochberg correction). Genes were filtered at adjusted p-value<0.05 and ranked by log₂ fold change. (B) Bean plots of log2CPM expression for ten representative genes in female and male LC cells. Median values are displayed as a long line and individual donor values are shown as short lines. Dotted line indicated overall mean. (C-F) Bubble plots for GO Biological processes (C-D), and Molecular functions (E-F) are displayed, with those up in male shown on the left (C,E) and down in male on the right (D,F). Bubble size indicates gene count, and color shows the −log10(adjusted p-value).

### Male Breast Shows Reduced Expression of Inflammatory Ligand Signatures and an Elevated Estradiol Gene Set Score in LC Cells

To better characterize the sex-specific signaling programs of male LC cells, we next assessed ligand-receptor interactions based on the combination of sex-specific downstream targets, expression of ligand by at least one cell type in the tissue, and expression of receptor in the LC population. This revealed a broad suppression of cytokine and immune signaling in luminal cells (Figure 4A), despite many of these having continued expression of the ligands across mammary cell populations (Figure 4B). This may be either due to a suppressed response, supported by significantly elevated expression of the cytokine signal suppressors SOCS2 and SOCS4 in male LC (data not shown), or due the decreased epithelial cell numbers resulting in lower local concentrations of these factors. These differences in cytokine activities are consistent with the GO analysis in Figure 3 suggesting generally decreased immune and signaling activities in male breast tissue. Amongst all of these, only the enrichment score for estradiol-responsive genes was higher in male LC than in female (Figure 4D). This observation is consistent with the higher *ESR1* mRNA observed in males than in females (adjusted p=0.0036; Figure 4F). *PGR* was also significantly elevated in male LC (adjusted p=0.039), and AR showed a non-significant trend to higher expression in male (adjusted p=0.09). *ESR2* did not differ significantly between sexes (adjusted p=0.46. Expression of hormone-metabolizing enzymes including *HSD17B1* (Figure 4E), *HSD17B12*, *CYP17A1*, *PTGES*, and *ALDH1A3,* also varied between male and female cell types, though whether they affect local estrogen concentrations, cannot be determine from mRNA expression data alone. HSD17B1 and HSD17B12, which convert estrone to estradiol, were enriched in male LC (Figure 4E), raising the possibility of autocrine estradiol production within this compartment. In male tissues, estrone arises predominantly from CYP19A1-mediated aromatization of circulating androgens in the stromal compartment,^32^ positioning epithelial HSD17B1/12 activity to locally amplify estradiol availability. Despite low circulating estrone in healthy men, epithelial HSD17B1-mediated intracrine conversion of estrone to estradiol may sustain local ER signaling, analogous to postmenopausal breast tissue.^33,34^ The clinical relevance of local estrogen biosynthesis in the male breast is further suggested by the high rates of gynaecomastia and male breast cancer observed in men receiving androgen deprivation therapy for prostate cancer, in whom suppression of testicular androgen production alters the androgen-to-estrogen ratio and unmasks estrogen-driven epithelial responses.^35^ The elevated *ESR1* and *PGR* mRNA in male LC is consistent with the molecular phenotype of ER⁺ breast tumors, and raises the question of whether LC may represent the normal epithelial population most relevant to male breast cancer.

**Figure 4.**
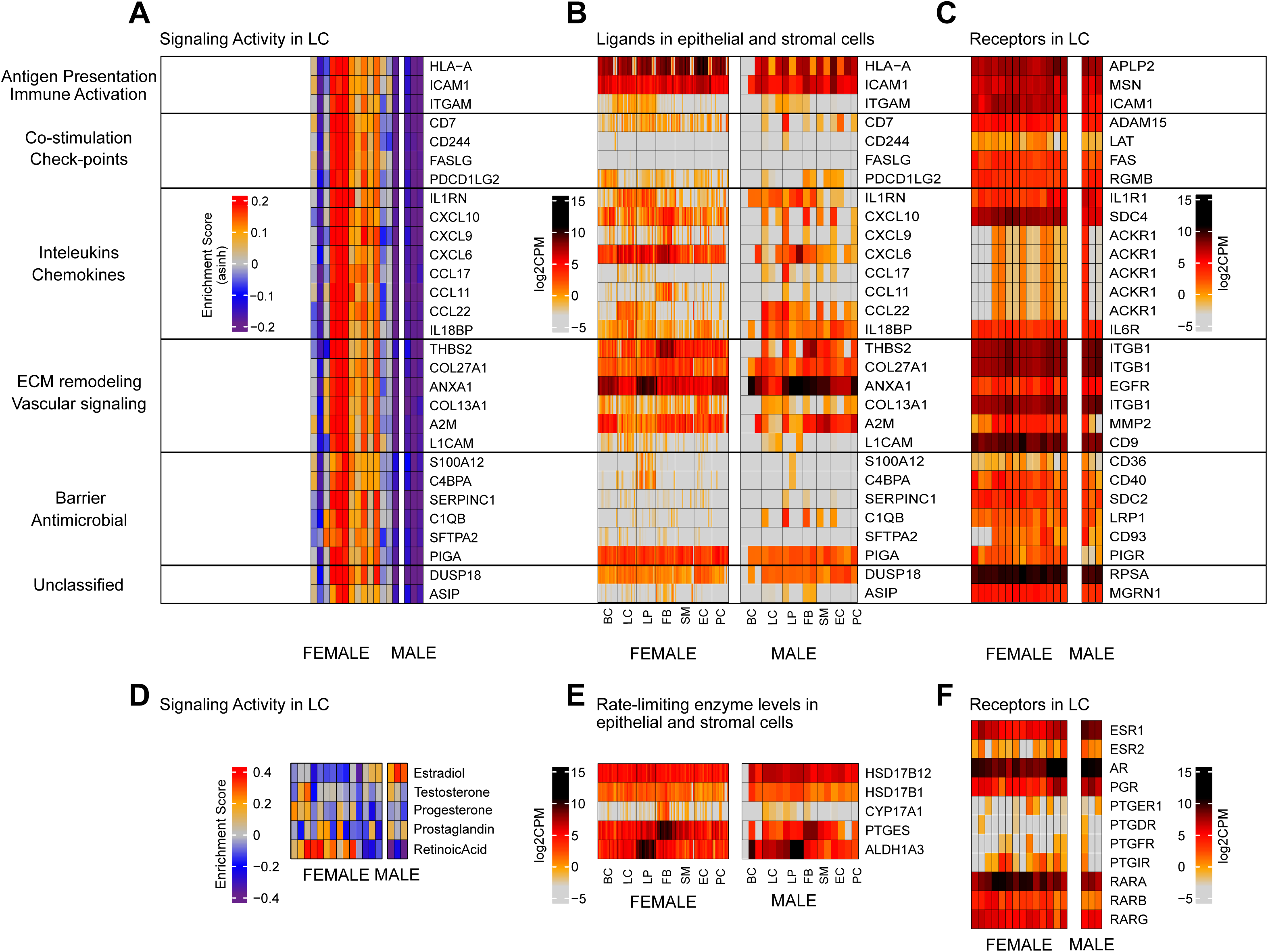
Male breast LC show decreased cytokine signaling but higher estradiol gene set enrichment scores compared with female LC cells. (A) Heatmap of per-sample GSVA enrichment scores for ligand-associated downstream regulons derived from NicheNet ligand-target predictions and MSigDB C2 curated gene sets for hormone responses. GSVA was computed on voom-normalized log2CPM expression values. Statistical significance between male and female LC was assessed using CAMERA competitive gene set testing (trend-adjusted variance, inter-gene correlation accounted for via allow.neg.cor=TRUE), with p-values adjusted by the Benjamini-Hochberg method. Only gene sets significant at FDR<0.05 are shown (n=49), all of which were downregulated in male LC with the exception of Estradiol which was upregulated (see D). (B) Heatmaps of mean log2CPM-normalized expression of the ligand (or rate limiting enzyme) associated with the top differentially expressed ligand regulons from (A) are shown for each donor across epithelial and stromal cell types (BC, LC, LP, FB, SM, EC, PC) in female and male. (C) Heatmap of mean log2CPM-normalized expression of LC receptors for the ligands shown in (B), for female and male donors separately. (D) Heatmap of GSVA enrichment scores for indicated steroid hormones for each sample. (E,F) Heatmap of mean log2CPM-normalized expression of (E) hormone-metabolizing enzymes (HSD17B12, HSD17B1, CYP17A1, PTGES, ALDH1A3) and (F) steroid hormone receptors (*ESR1*, *ESR2*, *AR*, *PGR*), prostaglandin receptors (*PTGER1*, *PTGDR*, *PTGFR*, *PTGIR*), and retinoic acid receptors (*RARA*, *RARB*, *RARG*) in LC across donors stratified by sex.

### Transcription Factor Activity Scores Differ Between Male and Female Breast Epithelial Cells

Inferred transcription factor activity scores, estimated by regulon-based enrichment analysis, showed sex-associated differences across the patient cohort (Figure 5). Unlike cytokine signaling activity which was strongly decreased across most ligands in male LC (Figure 4A, 3D,F), transcription factors showed a mixture pattern, with some showing elevated activity in males while others were suppressed (Figure 5). Overall, the differential expression of these factors was congruent with the previous ligand-receptor and gene ontology analyses. Consistent with these previous analyses, we observed that male LC exhibited decreased signs of inflammatory cytokine/growth factor activity (STAT3, NFKB1/2, REL, IRF5, ATF2/4, AP1, FOS/JUN/JUND, Figure 5). Consistent with the elevated ribosomal RNA production, UBTF, a master regulator of rRNA transcription, was elevated in male LCs. This alongside elevated PPARA and NRF1 fits with a metabolically active state. Male LC also showed higher inferred activity of chromatin regulators including SMARCA5, CTCFL, and HDAC1, alongside elevated enrichment scores for transcription factors with established roles in positional patterning and epithelial morphogenesis, among them HOX family members HOXA10, HOXB7, HOXC6,^36^ HOX cofactor MEIS1,^37^ homeodomain factor MSX2,^38^ zinc-finger regulator FEZF1, T-box factor TBX3,^39^ lymphatic and epithelial identity regulator PROX1,^40^ and luminal alveolar transcription factor ELF5.^41^ Several of these factors are known to be active during early mammary gland development and are progressively modulated as the gland undergoes hormone-driven morphogenetic remodeling postnatally. Their relative enrichment in male LC is consistent with the comparatively limited postnatal remodeling of the male mammary gland, though whether this reflects a developmental or a sex-specific transcriptional feature of adult male LC cell identity remains to be determined.

**Figure 5.**
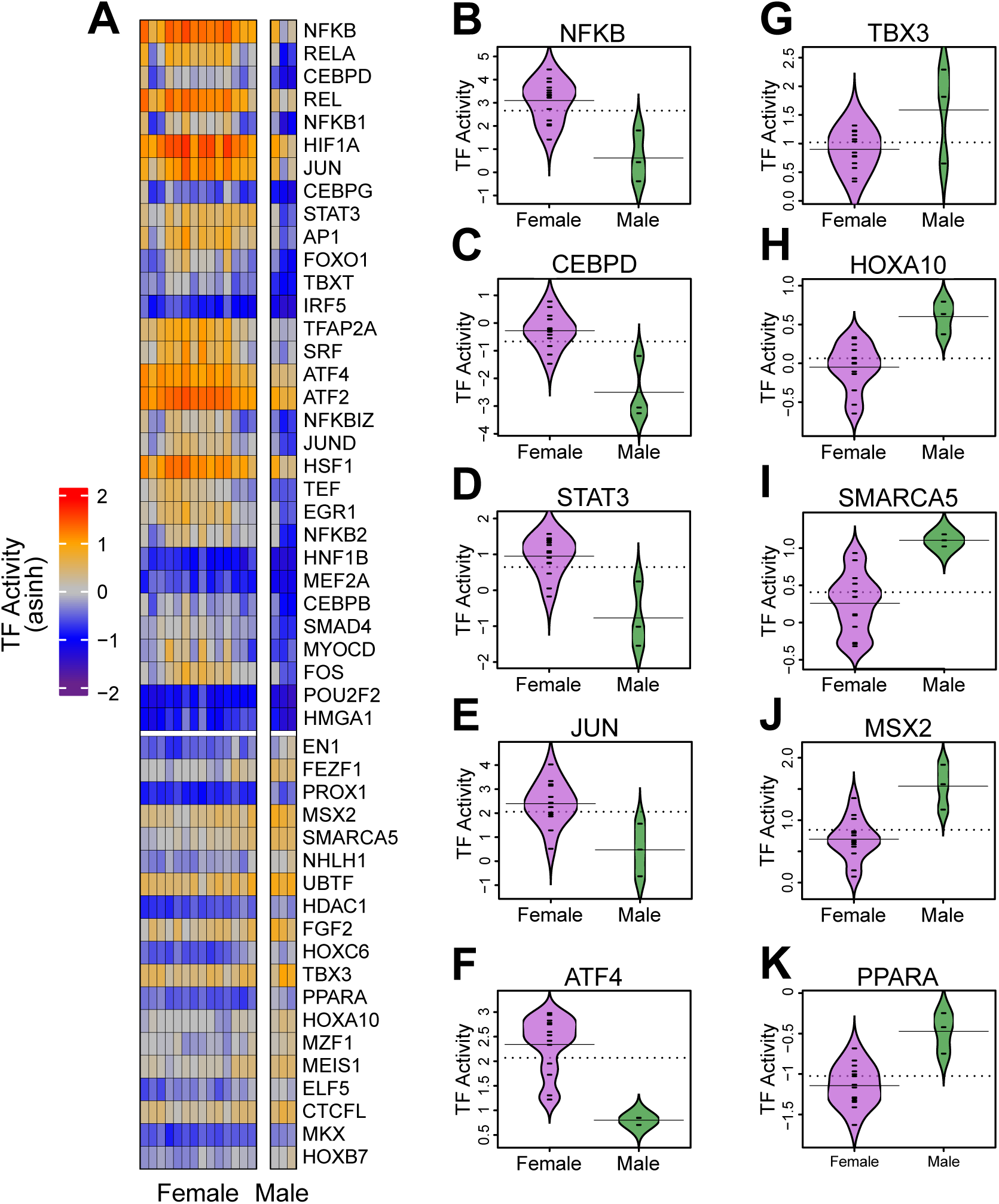
Sex-associated differences in inferred transcription factor activity scores in breast epithelial cells. (A) Heatmap of the top 49 TFs with significantly differential activity between male and female LC (asinh(score/2)-transformed) across donors stratified by sex (FDR<0.05). Activity scores were estimated using the Univariate Linear Model (ULM, decoupleR) applied to voom-normalized log2CPM values with the CollecTRI regulon database (minimum 5 target genes per TF), and differential activity was assessed by limma with Empirical Bayes moderation and Benjamini-Hochberg correction. (B–K) Beanplots comparing inferred ULM activity scores for ten TFs selected as the top 5 female-enriched and top 5 male-enriched by log₂ fold change (female in purple and male in green): (B) *NFΚB* (female-enriched); (C) *CEBPD* (female-enriched); (D) *STAT3* (female-enriched); (E) *JUN* (female-enriched); (F) *ATF4* (female-enriched); (G) *TBX3* (male-enriched); (H) *HOXA10* (male-enriched); (I) *SMARCA5* (male-enriched); (J) *MSX2* (male-enriched); (K) *PPARA* (male-enriched). Donor-level values and medians are shown. TF annotations describe known biological roles; these are provided for interpretive context and do not imply that the activity differences shown here have been validated in male breast tissue.

## DISCUSSION

The normal adult male breast has not previously been characterized at single-cell resolution, and the dataset presented here provides that foundation. Across male donors, the mammary epithelium, although was heterogeneous, was markedly poor in basal and luminal progenitor cells and instead comprised predominantly LC-state. This compositional pattern was distinct from any female donor and independently supported by transcriptomic analysis and by the phenotypically defined luminal-like compartment across the broader eight-donor flow cytometric dataset, the gate least susceptible to stromal contamination in male tissue. Male LC were further distinguished from their female counterparts by elevated ESR1 and PGR mRNA, enrichment of RNA processing and ribosome biogenesis gene programs, reduced expression of genes associated with inflammatory cytokine and growth factor signaling, elevated estradiol gene set enrichment scores, and higher inferred activity of transcription factors associated with positional patterning and epithelial morphogenesis. That these differences are recovered consistently across differential expression, gene ontology enrichment, ligand profiling, and regulon-based transcription factor inference, four analytically independent approaches applied to the same dataset, strengthens confidence in their biological reality and provides the first reproducible transcriptional description of the normal male LC cell.

Considered together, these properties suggest that adult male LC occupy a metabolically competent but mitotically restrained state, biosynthetically poised rather than quiescent in the strict sense, and distinct from cellular senescence, which is characterized by irreversible growth arrest and a pro-inflammatory secretory phenotype directly contrary to the suppressed cytokine signaling observed here. The elevation of RNA processing and ribosomal gene programs indicates retained translational capacity, consistent with the ability of male-derived cells to proliferate in culture and with the clinical demonstration that male mammary epithelium retains the capacity for hormone-driven expansion in gynecomastia.^4,42,43^ At the same time, the suppression of inflammatory cytokine and growth factor signaling programs observed independently across gene ontology, ligand expression, and downstream transcription factor analyses is coherent with the in vivo mitotic restraint of the male epithelium and with the slower cycling of male-derived cells observed under culture conditions that support rapid proliferation of equivalent female populations. The over-expression of SOCS2 and SOCS4 in male LC (data not shown) raises the possibility that this suppression is at least partly cell-intrinsic, reflecting a differential transcriptional response to cytokine exposure rather than simply a consequence of reduced local factor concentrations in the more sparsely populated male epithelium though both mechanisms may contribute. Direct measurement of cytokine signal transduction downstream of defined stimuli, as previously performed in female breast populations,^44^ would be needed to resolve this distinction. Notably, a recent study demonstrated that exogenous androgen therapy in individuals recorded female-at-birth induces broad transcriptional and translational silencing in luminal breast cells alongside reduced immune signaling at the tissue level, mediated in part through paracrine mechanisms downstream of androgen-receptor activation in hormone-responsive cells.^45^ The convergence of these androgen-induced changes with the transcriptional properties we observe in the male LC of cisgender men suggests that some features of the male LC transcriptional state reflect the cumulative effects of lifelong androgen exposure rather than acute hormonal signaling, though the developmental and induced components of this state cannot be fully disentangled in cross-sectional data from either study.

The simultaneous enrichment of ESR1, PGR, and the estrogen-metabolizing enzymes HSD17B1 and HSD17B12 in male LC raises the further question of whether autocrine estradiol production contributes to the maintenance of this population in the adult male gland, an observation that motivates, but does not establish, a functional role and one that will require targeted metabolic and signaling studies to evaluate. The observation that androgen therapy in the female breast suppresses PGR while leaving ESR1 accessibility largely intact ^45^ is notable in the context of our finding that both *ESR1* and *PGR* are elevated in LC of men. This divergence suggests that the hormone receptor profile of the developmentally male breast is not simply a consequence of androgen-mediated suppression of the female receptor program but may reflect a distinct transcriptional state established during development, a distinction that prospective functional studies will need to address. It should be noted that the male and female donors contributing scRNA-seq data differ in mean age, male scRNA-seq donors mean age 66 years, female scRNA-seq donors mean age 41 years, and age-related changes in hormone receptor expression, including the well-documented decline in PGR with advancing age, represent a confounding variable that cannot be fully resolved from the present cross-sectional dataset; prospectively collected age-matched donors will be required to isolate the contribution of biological sex from that of age to the hormone receptor differences reported here.

The higher inferred activity of developmental patterning regulators, HOX family members, MEIS1, MSX2, TBX3, ELF5, and associated chromatin factors including SMARCA5, CTCFL, and HDAC1, in cells that have undergone comparatively limited postnatal morphogenetic remodeling is consistent with the interpretation that male LC retain aspects of an earlier transcriptional-state. Whether this constitutes a sex-specific feature of male LC identity or a reflection of incomplete developmental progression cannot be determined from the present data. Several of these factors carry established prognostic associations in female breast cancer that are germane to this interpretation: TBX3 overexpression predicts poor survival specifically in luminal A tumors,^39,46^ elevated ELF5 predicts shortened metastasis-free and overall survival in the same subtype,^47,48^ and SMARCA5 overexpression correlates with advanced stage disease and reduced overall survival.^49^ Conversely, MEIS1 functions as a growth suppressor that is downregulated in breast invasive carcinoma^50^ and MSX2 expression associates with ER positivity and prolonged patient survival,^51^ suggesting that the patterning program of male LC does not map onto a uniformly aggressive transcriptional state but rather reflects a complex regulatory landscape whose functional consequences in the male breast remain to be determined.

To assess whether the transcriptional identities of male mammary epithelial populations bear systematic relationships to breast cancer molecular subtypes, we correlated gene signatures derived from male LC and LP independently with subtype-specific expression profiles across TCGA breast cancer samples. The two signatures showed strikingly opposing patterns of subtype association. The male LC signature correlated most strongly and positively with Luminal A tumors and Normal-like breast cancer, with progressively weaker association in HER2-enriched tumors and significant negative correlations with Basal-like and Luminal B subtypes. The male LP signature showed the precise inverse, strongest positive correlation with Basal-like tumors and significant negative correlations with Luminal A disease. This reciprocal pattern recapitulates in the male breast the cell-of-origin associations previously established for female mammary epithelial populations, where LP are transcriptionally proximate to basal-like and BRCA1-associated cancers and LC to ER-positive luminal disease,^19^ and demonstrates that these relationships are conserved across sexes despite the marked compositional and transcriptional differences between male and female epithelium documented here. The strong LC-Luminal A correlation warrants reconciliation with the clinical evidence that male ER⁺ tumors behave more aggressively than their female counterparts. A substantial component of this excess mortality is attributable to later stage at diagnosis rather than intrinsic tumor biology, and stage-adjusted survival differences between male and female ER⁺ disease are considerably attenuated in population-based analyses.^52^ Transcriptional similarity between normal cells and tumor subtypes reflects differentiation state and likely cell of origin, rather than tumor behavior. Determinants of aggressiveness in male breast cancer will require direct comparison of normal male LC cells with matched tumor datasets, which this atlas now enables.

These observations also invite consideration of their relevance to male breast cancer, though several points of precision are warranted before drawing conclusions. Independent studies have shown that normal hormone-responsive LC correlate most strongly with PAM50 luminal A and B breast cancer classifications, and that androgen therapy reduces these correlations, consistent with the interpretation that the hormone-sensing luminal compartment is the normal epithelial population most transcriptionally proximate to luminal breast cancers, irrespective of the sex of the donor.^19,45^ Transcriptional similarity between a normal epithelial population and a tumor type does not establish a cell-of-origin relationship, a transformed LP or basal cell can acquire a luminal transcriptional profile under oncogenic conditions, and the present data cannot exclude this possibility.^53^ It should also be noted that some male donors not included in the sequenced cohort showed higher proportions of cells within the phenotypically defined basal-like compartment by flow cytometry, and that at the transcriptional level, a stromal subpopulation in male tissue shared expression features with BC, distinguished only by differential KRT5/KRT14 expression in true BC versus RGS5-expressing cells in the stromal subpopulation. This observation introduces interpretive caution regarding the completeness of basal cell depletion in the male breast, while simultaneously reinforcing the need for transcriptomic rather than purely phenotypic classification of male mammary cell populations. With these qualifications stated, the convergence of three independent observations is noteworthy: i) LC are numerically dominant in the male epithelium, making them the population with the greatest prior probability of being the cell of origin for male mammary tumors, acknowledging that the epithelial composition of men who subsequently develop breast cancer may differ from the general male population characterized here, a question that prospective longitudinal studies will be required to address; ii) ESR1 and PGR mRNA are significantly elevated in male LC relative to all other male epithelial populations, concordant with the high rates of ER and PR positivity that characterize MBC across population-based series;^54^ and the developmental patterning programs enriched in male LC include factors associated with poor prognosis in female luminal breast cancer.^39,46–49^ iii) None of these observations establishes causation, but together they define male LC as the most biologically plausible normal cell population in which to investigate the origins of MBC. A particularly informative comparison will be between MBC and postmenopausal female luminal tumors, two ER⁺ disease contexts with similar current hormonal environments but divergent lifetime exposure histories, which may help delineate the contributions of hormonal history from those of sex-specific epithelial cell biology. Testing the cell-of-origin question directly will require transformation assays, patient-derived models, and systematic comparison of normal male LC transcriptomes with MBC tumor datasets.

Our male cohort had eight donors characterized for epithelial content and subtype composition, a cohort size that itself reflects a fundamental biological constraint of the male breast as a study system. Viable EpCAM⁺ epithelial populations were detected by FACS in 75% of male donors processed, compared with all female donors processed in parallel, and only three donors yielded sufficient viable cell numbers for scRNA-seq. This attrition rate is documented here as biologically meaningful information for future study design; it also means that male breast scRNA-seq datasets of the scale routinely achieved in female studies are not currently obtainable. Future studies could incorporate cell multiplexing strategies^55–57^ to pool cells from multiple male donors prior to library preparation, enabling characterization of a broader donor range.

Age-associated transcriptional differences cannot be excluded as a partial contributor to the observed signals. This confound cannot be resolved from the current cross-sectional dataset and will require prospectively collected, age-matched donors to address definitively. Circulating hormone levels were not systematically captured at the time of tissue collection; prospective studies incorporating paired serum hormone measurements would allow direct correlation of systemic hormonal context with LC cell transcriptional state and help distinguish cell-autonomous programs from those driven by circulating hormone concentrations. Finally, protein-level validation of key findings including ESR1 and PGR expression, functional interrogation of male LC cell hormone responsiveness, and direct transcriptomic comparison with male breast tumor datasets^8,58–61^ represent the now tractable next steps made possible with this male single cell atlas.

This dataset establishes the first cellular and transcriptional reference for the normal human male breast. It documents that the male mammary epithelium is compositionally and transcriptionally distinctive, characterized by LC dominance, elevated hormone receptor expression, biosynthetic competence, suppressed cytokine signaling, and persistent developmental patterning programs. These properties were reproduced across donors and across independent analytical methods. Similarly, this study identifies important logistic considerations in the collection and purification of mammary cells in male which could help inform future experimental designs. The atlas, differential expression data, and transcriptional characterization of male LC cell identity that this study provides, constitute a resource for prospective functional investigation, allowing the contextualization of germline susceptibility variants in their normal cellular environment, and for interrogating the biology of a breast cancer subtype whose cellular origins have remained inaccessible in the absence of a defined normal tissue reference.

## METHODS

### Male and female breast specimen

Normal human mammary tissues were collected under an IRB-approved Mayo Clinic enterprise-wide protocol (N.K.) through the Regenerative Breast Biobank at Mayo Clinic Rochester, with two additional samples contributed from the breast tissue collection of D.R. at Mayo Clinic Jacksonville under a compatible IRB protocol. All samples were processed using identical procedures in N.K.’s laboratory at Mayo Clinic Rochester, ensuring methodological consistency across collection sites. Written informed consent was obtained from all living donors; post-mortem samples were obtained under Minnesota Anatomical Gift Authorization. The cohort comprised 8 male donors and 27 female donors from elective surgical resection or post-mortem donation. Detailed donor metadata are available from the corresponding author (N.K.) upon reasonable request and will be included in the final version of this manuscript.

### Tissue dissociation and fluorescence-activated cell sorting

Cryopreserved or fresh tissue organoids were dissociated sequentially with Trypsin-EDTA (1–5 mL), Dispase (2 mL), and DNase I (200 μL) in HBSS + 2% FBS (HF), filtered through a 40-μm cell strainer, and centrifuged at 1,200 rpm, 4°C. Viable cells were stained with DAPI and an antibody cocktail (1:50 each: anti-CD31, anti-CD34, anti-CD45, anti-EpCAM, anti-CD49f) for 30 minutes on ice. FACS sorting was performed on a BD FACSMelody. Viable Lin⁻ (CD31⁻CD34⁻CD45⁻) DAPI⁻ cells were sorted into luminal-like (EpCAM^hi^CD49f^low^), luminal progenitor-like (EpCAM⁺CD49f⁺), basal-like (EpCAM^low^CD49f^hi^), and stromal-like (EpCAM^low^CD49f^low^) fractions for 2D expansion. Viable (DAPI-) cells were sorted and maintained as single cell suspensions for scRNA-Seq experiments.

### 2D proliferation and 3D organoid assays

FACS-sorted cells were expanded in DMEM/F12 complete medium, containing 5% FBS, 10 ng/mL cholera toxin, 0.5 μg/mL hydrocortisone, 1 μg/mL insulin, 10 ng/mL EGF and 1% penicillin/streptomycin, in collagen-coated dishes. Proliferation was assessed by live-cell imaging (IncuCyte S3 2018C) of 10,000 cells per well over 116 hours. Doubling times were calculated from phase-object confluence curves. For 3D assays, 500–5,000 cells were embedded in Matrigel (50 μL per chamber), solidified at 37°C, and cultured in complete medium. Brightfield images were acquired on a Cytation5.

### Single-cell RNA sequencing library preparation

FACS-sorted cells were processed on the 10x Genomics Chromium Controller using the Next GEM Single Cell 3′ v3.1 Dual Index kit, as previously published.^62,63^ Libraries were assessed by Agilent Bioanalyzer and sequenced as PE150 reads on an Illumina NovaSeq 6000.

### Read processing and quality control

FASTQ files were aligned to GRCh38 (v2020-A) using Cell Ranger v6.1.2. Raw count matrices were imported into Seurat and duplicate gene names were collapsed. Metadata describing donor, sex, mutation status and sequencing chemistry were added. In parallel, public scRNA–seq data from non-cancerous female donors sequenced^7^ were downloaded as count matrices and loaded into Seurat with minimal filtering (cells with ≥200 detected genes and genes expressed in ≥3 cells) and annotated with sample metadata. Per–cell metrics including the number of detected genes (nFeature_RNA), number of total counts (nCount_RNA) and the fraction of mitochondrial reads (percent.mt) were calculated. Cells with extremely low gene counts, unusually high UMI counts, or elevated mitochondrial content (>3 median absolute deviations from the median as determined using scater (1.28.0,^64^) were removed. Genes detected in fewer than five cells were discarded.

Counts were log-normalized per cell and the top 2000 highly variable features identified per sample using Seurat’s (5.4.0) FindVariableFeatures with the vst method.^65^) Data were scaled and centered using ScaleData, regressing out nCount_RNA and percent.mt to mitigate technical confounders. Principal–component analysis (PCA) was performed on the scaled variable features. Clusters were then identified using the Louvain algorithm implemented in Seurat’s FindClusters with resolution parameters of 0.5 for the Mayo Clinic samples and 0.8 for the public available samples from Nee et al. These resolutions provided clusters corresponding to known mammary epithelial, stromal and immune lineages. Droplet scRNA–seq data can contain droplets encapsulating two or more cells (doublets). Such doublets often cluster between real cell types and can confound downstream analysis.^66^ We used DoubletFinder on the pre–processed Seurat objects to identify and remove doublets. The expected doublet rate was estimated at ∼5% based on 10x Genomics chemistry, and the homotypic doublet rate was corrected using cell type proportions. The pK parameter, representing the number of principal components used to identify artificial doublets, was determined by maximizing the mean–variance normalized bimodality coefficient across a sweep of values as recommended.^66^ Cells predicted as doublets were excluded from further analysis.

### Imputation, batch correction, and cell type labelling

To alleviate sparsity due to dropout, we applied the Adaptively thresholded Low–Rank Approximation (ALRA) algorithm implemented in SeuratWrappers. ALRA assumes that the true expression matrix is non–negative, low–rank and contains many biological zeros. It first computes a low–rank approximation of the observed matrix by singular–value decomposition, with the rank automatically selected, and then thresholds each gene’s values by the magnitude of the most negative entry to restore biological zeros.^67^ This procedure selectively imputes technical zeros while preserving genuine zero expression.^67^ After imputation, counts were projected into a batch–invariant latent space using the deep–learning–based SCALEX algorithm. SCALEX learns a generalized encoder that disentangles batch–specific components and maps cells into a common cell–embedding space, allowing online integration of new datasets without retraining and outperforming other integration methods on partially overlapping data.^68^ The latent features returned by SCALEX were imported back into R for downstream analysis. A shared nearest-neighbor graph was then constructed from the latent space and clusters were identified using Seurat’s Louvain algorithm (resolution 0.5).

### Visualization and cluster annotation

Cell-type identities in the integrated Mayo Clinic reference dataset were assigned using canonical lineage marker genes. After integration and clustering, expression of known mammary epithelial, stromal, endothelial, and immune markers was examined across the UMAP embedding to determine the biological identity of each cluster. Gene expression was visualized using custom density-based visualization functions that projected expression values onto the UMAP embedding using hexagonal binning to reduce overplotting. Panels of lineage markers were examined iteratively to identify clusters enriched for specific cellular programs. These included luminal epithelial markers (EPCAM, KRT8, KRT18), BC markers (KRT5, KRT14, ACTA2, ACTG2, MYLK, TAGLN, KRT17), LP markers (PROM1, KIT, ELF5, ALDH1A3, S100A8, S100A9), LC markers (ANKRD30A, GATA3, HSPB1), endothelial markers (PECAM1, CDH5, ENG, FLT1), stromal and fibroblast markers (PDGFRA, COL1A2, FN1, THY1, DCN, LUM), and immune markers (PTPRC, CD68, CSF1R, CD14). Clusters were assigned to mammary lineages based on the enrichment and spatial localization of these marker genes within the UMAP embedding, and the resulting lineage labels were stored in the Seurat metadata. This annotated dataset was subsequently used as the reference for cell-type label transfer to the external dataset.

The manually annotated Mayo Clinic reference dataset was then used to assign cell-type identities to the remaining samples from Nee et al. using Seurat’s anchor-based label transfer framework.^65^ Transfer anchors between the annotated reference object and each query dataset were then identified using the FindTransferAnchors function with the first thirty principal components. These anchors represent pairs of transcriptionally similar cells between the reference and query datasets and provide a mapping between the two datasets in a shared feature space. Cell-type labels from the reference object were subsequently projected onto the query cells using Seurat’s TransferData function, which assigns the most probable cell identity to each cell based on the anchor relationships. The predicted labels were stored in the Seurat metadata and used as the primary cell-type annotations for downstream analyses. Prediction scores generated by the transfer procedure were used to assess assignment confidence, and cells with low prediction scores (<0.5) were flagged for manual review. Marker gene expression patterns were examined in these cases to confirm or refine lineage assignments prior to downstream analyses.

### Cell-type composition analysis

To compare the relative abundance of mammary epithelial lineages across donors and sexes, the distribution of annotated cell types was quantified for each sample. Cells were grouped by donor and lineage annotation, and the counts of basal (BC), luminal progenitor (LP), and differentiated luminal (LC) epithelial cells were computed for each donor. These counts were converted to proportions relative to the total number of epithelial cells per donor to account for differences in cell recovery across samples. The resulting compositional profiles were visualized using ternary plots, which represent the relative contributions of BC, LP, and LC populations within each donor sample. Visualization was performed using the ‘ggtern’ package (4.0.0) in R, with samples colored and shaped according to sex. This representation allowed simultaneous visualization of the balance between epithelial lineages within each donor and facilitated comparison of lineage composition between male and female tissues. By projecting each donor onto the ternary coordinate system defined by the three epithelial states, shifts in lineage proportions could be readily assessed while preserving the constraint that the three components sum to one.

### Pseudobulk aggregation and differential gene expression

To compare transcriptional programs between sexes while accounting for donor-level variability and avoiding skewing and over-inflation of significance due to large or variable cell numbers, differential expression analyses were performed using a pseudobulk framework. For each annotated lineage, raw counts from all cells belonging to the same donor were summed using Seurat’s AggregateExpression function, producing a donor-level expression matrix in which each column represented the aggregated transcriptome of a given cell type within a donor. Only donors contributing cells to a given lineage were included in that lineage-specific analysis. The resulting pseudobulk count matrices were analyzed using the edgeR–limma workflow. Briefly, counts were imported into edgeR (3.42.4) and normalized for library size using trimmed mean of M-values normalization. The normalized counts were then transformed using the voom method, which models the mean–variance relationship of RNA-seq data and assigns precision weights to each observation.^69^ Linear models were fitted using limma (3.56.2) with sex as the primary variable of interest, and empirical Bayes moderation was applied to stabilize variance estimates across genes. Differentially expressed genes were identified based on moderated statistics and false discovery rate correction using the Benjamini–Hochberg method.

### Gene ontology enrichment analysis

Functional interpretation of differentially expressed genes identified in the pseudobulk analyses was performed using Gene Ontology (GO) enrichment analysis. Differential gene expression between male and female donors was first identified using the pseudobulk limma-voom framework described above. Genes showing significant differential expression following false discovery rate correction were separated into up-regulated and down-regulated gene sets. Enrichment for GO biological process terms was then assessed using the goana function from the limma package, which tests whether genes associated with a given ontology category are overrepresented among the differentially expressed genes relative to the background of all expressed genes. The analysis accounts for gene length and other potential biases inherent to RNA-seq data. Significance values were adjusted for multiple testing using the Benjamini–Hochberg procedure. Enriched GO categories were summarized by ranking terms according to statistical significance and visualized using bar and dot plots showing the proportion of genes within each category that were differentially expressed. This analysis provided functional context for the transcriptional differences observed between sexes and highlighted biological processes associated with sex-specific transcriptional programs within individual mammary epithelial lineages. To ensure that GO enrichment results were not driven by Y-chromosome gene dosage, the analysis was repeated after excluding all Y-linked genes (*RPS4Y1*, *EIF1AY*, *DDX3Y*, *PCDH11Y*, *SPRY3*) from both the DE gene list and the background universe; results were identical, confirming that the enriched terms are not an artifact of Y-chromosome gene dosage.

### Transcription-factor activity inference

To infer upstream regulatory programs associated with the observed transcriptional differences, transcription-factor (TF) activity scores were computed from the pseudobulk expression matrices using decoupleR (2.6.0s). The analysis used the human CollecTRI regulatory network, which integrates experimentally supported TF–target interactions and encodes the direction of regulation for each interaction.^69^ TF activities were estimated using the univariate linear model (ULM) method implemented in run_ulm, which evaluates the association between expression of a regulator’s target genes and the regulator’s signed interaction weights. Only regulators represented by at least five target genes in the expression matrix were retained. The resulting TF activity scores represent donor-level estimates of regulatory activity rather than gene expression. Differential TF activity between male and female donors was then assessed using the same limma modeling framework applied to gene expression data. This approach enabled identification of transcriptional regulators whose inferred activity differed between conditions while accounting for donor-level replication.

### Ligand–receptor signaling pathway analysis

To investigate intercellular signaling programs associated with sex differences in mammary tissue, ligand-centered signaling pathways were defined using the NicheNet framework^70^ which integrates experimentally supported ligand–receptor interactions and downstream ligand–target regulatory relationships. For five endocrine ligands (testosterone, progesterone, prostaglandin E2, retinoic acid, and estradiol), gene sets were instead derived from curated transcriptional response signatures in MSigDB (C2 collection). Candidate ligands were identified from genes expressed in potential sender cell populations, including basal epithelial cells, luminal progenitors, fibroblasts, smooth mucscle cells, endothelial cells, pericytes, and immune cells. For each ligand, a discrete gene list of targets expected to be activated downstream of that ligand in the receiver cells was created using the make_discrete_ligand_target_matrix from the R package nichenetr (2.2.1.1) based on the ligand-target database from NicheNet v2.

Pathway activity scores were then calculated using Gene Set Variation Analysis (GSVA), which computes a non-parametric enrichment score for each pathway in each sample based on coordinated expression of the pathway genes.^71^ GSVA was applied to the voom log-CPM expression matrix to generate enrichment scores for each ligand-associated signaling program across samples. Gene sets with fewer than three genes detected in the expression matrix were excluded prior to testing. To identify pathways showing significant sex-associated activity differences, competitive gene-set testing was performed using the camera method implemented in limma^72^, which evaluates whether genes within a pathway exhibit systematic differential expression relative to the background while accounting for inter-gene correlation. Resulting enrichment scores and ligand expression patterns across cell types were visualized to characterize signaling programs differentially active between male and female mammary tissues.

## DATA AVAILABILITY

Male scRNA-seq datasets and extensive supporting analyses generated for this study are available upon reasonable request and will be made publicly available upon acceptance.

## ACKNOWLEDGEMENTS

This work was supported in part by Mayo Clinic–National Cancer Institute (NCI) SPORE in Breast Cancer (P50 CA116201-12CEP) and Ovarian Cancer (P50 CA136393-11CEP) Career Enhancement Awards to N.K. M.R. was supported by the National Institute of General Medical Sciences of the National Institutes of Health T32 training grant (T32-GM144233). A.J. was supported by the Mayo Clinic Summer Undergraduate Research Fellowship. M-I.I.R. was supported by Cancer Research Society in partnership with Canadian Institutes of Health Research (CIHR), Institute of Cancer Research (ICR) and IRIC. D.J.H.F.K. was supported by philanthropic funds from the Marcelle and Jean Coutu Foundation through the IRIC and held a Junior 1 Chercheurs-boursiers salary award from the Fonds de recherche du Québec, Santé (FRQS; #283502). F.J.C. was supported by NCI grants R35 CA253187 and P50 CA116201 and by the Breast Cancer Research Foundation. The authors thank the donors for contributing tissue samples to the Mayo Clinic Regenerative Breast Biobank.

## AUTHOR CONTRIBUTIONS

N.K.: Conceived and initiated the study; generated all primary data; secured funding; supervised the project; and co-wrote the manuscript. D.J.H.F.K.: Co-developed analytical approaches; led data analysis and visualization; supervised computational work; and co-wrote the manuscript. M.-I.I.R.: Performed data curation, formal analysis, and methodology development; contributed to writing. S.M.M.A., M.L.R., A.J.: Investigation, data analysis, and methodology. K.C.: Investigation. J.W.J., S.A.M., A.C.D., J.C.B.: Resources and manuscript review. D.R.: Provided additional unpublished scRNA-seq datasets and reviewed the manuscript. F.J.C., S.Y., A.S., C.P.E.: Conceptual input and manuscript review. M.E.S.: Provided critical feedback that guided additional data analysis and reviewed the manuscript.

